# Strategies for analyzing bisulfite sequencing data

**DOI:** 10.1101/109512

**Authors:** Katarzyna Wreczycka, Alexander Gosdschan, Dilmurat Yusuf, Björn Grüening, Yassen Assenov, Altuna Akalin

**Author notes:** Equal contributions (Katarzyna Wreczycka, Alexander Gosdschan).

## Abstract

DNA methylation is one of the main epigenetic modifications in the eukaryotic genome; it has been shown to play a role in cell-type specific regulation of gene expression, and therefore cell-type identity. Bisulfite sequencing is the gold-standard for measuring methylation over the genomes of interest. Here, we review several techniques used for the analysis of high-throughput bisulfite sequencing. We introduce specialized short-read alignment techniques as well as pre/post-alignment quality check methods to ensure data quality. Furthermore, we discuss subsequent analysis steps after alignment. We introduce various differential methylation methods and compare their performance using simulated and real bisulfite sequencing datasets. We also discuss the methods used to segment methylomes in order to pinpoint regulatory regions. We introduce annotation methods that can be used for further classification of regions returned by segmentation and differential methylation methods. Finally, we review software packages that implement strategies to efficiently deal with large bisulfite sequencing datasets locally and we discuss online analysis workflows that do not require any prior programming skills. The analysis strategies described in this review will guide researchers at any level to the best practices of bisulfite sequencing analysis.

## 1. Introduction

Cytosine methylation (5-methylcytosine, 5mC) is one of the main covalent base modifications in eukaryotic genomes. It is involved in epigenetic regulation of gene expression in a cell-type specific manner. It is reversible and can remain stable through cell division. The classical understanding of DNA methylation is that it silences gene expression when occurs at a CpG rich promoter region [1]. It occurs predominantly on CpG dinucleotides and seldom on non-CpG bases in metazoan genomes. The non-CpG methylation has been mainly observed in human embryonic stem and neuronal cells [2],[3]. There are roughly 28 million CpGs in the human genome, 60–80% are generally methylated. Less than 10% of CpGs occur in CG-dense regions that are termed CpG islands in the human genome [4]. It has been demonstrated that DNA methylation is also not uniformly distributed over the genome, but rather is associated with CpG density. In vertebrate genomes, cytosine bases are usually unmethylated in CpG-rich regions such as CpG islands and tend to be methylated in CpG-deficient regions. Vertebrate genomes are largely CpG deficient except at CpG islands. Conversely, invertebrates such as *Drosophila melanogaster* and *Caenorhabditis elegans* do not exhibit cytosine methylation and consequently do not have CpG rich and poor regions but rather a steady CpG frequency over the genome [5]. DNA methylation is established by DNA methyltransferases DNMT3A and DNMT3B in combination with DNMT3L and maintained through/after cell division by the methyltransferase DNMT1 and associated proteins. DNMT3a and DNMT3b are in charge of the de novo methylation during early development. Loss of 5mC can be achieved passively by dilution during replication or exclusion of DNMT1 from the nucleus. Recent discoveries of ten-eleven translocation (TET) family of proteins and their ability to convert 5-methylcytosine (5mC) into 5-hydroxymethylcytosine (5hmC) in vertebrates provide a path for catalysed active DNA demethylation [6]. Iterative oxidations of 5hmC catalysed by TET result in 5-formylcytosine (5fC) and 5-carboxylcytosine (5caC). 5caC mark is excised from DNA by G/T mismatch-specific thymine-DNA glycosylase (TDG), which as a result returns cytosine residue back to its unmodified state [7]. Apart from these, mainly bacteria but possibly higher eukaryotes contain base modifications on bases other than cytosine, such as methylated adenine or guanine [8].

One of the most reliable and popular ways to measure DNA methylation is bisulfite sequencing. This method, and related ones, allow measurement of DNA methylation at the single nucleotide resolution. In this review, we describe strategies for analyzing data from bisulfite sequencing experiments. First, we introduce high-throughput sequencing techniques based on bisulfite treatment. Next, we summarize algorithms and tools for detecting differential methylation and methylation profile segmentation. Finally, we discuss management of large datasets and data analysis workflows with a guided user interface. The computational workflow summarizing all the necessary steps is shown in Figure 1.

**Figure 1.**
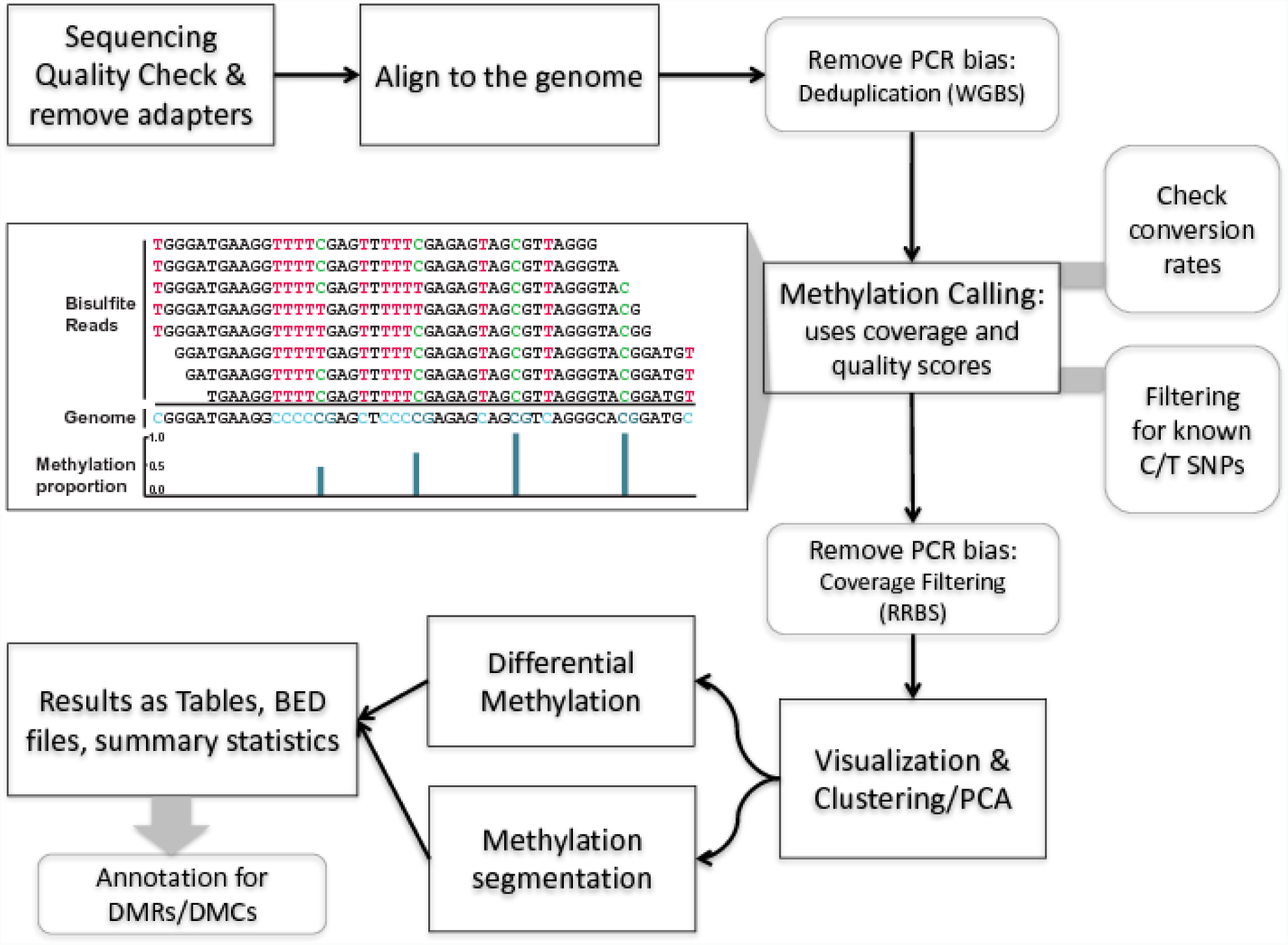
Workflow for analysis of DNA methylation using data from bisulfite sequencing experiments.

## 2. Bisulfite sequencing for detection of methylation and other base modifications

Techniques for profiling genome-wide DNA methylation fall into four categories: methods based on restriction enzymes sensitive to DNA methylation (such as MRE-seq), methylcytosine-specific antibodies (such as methylated DNA immunoprecipitation using MeDIP-seq [9]), methyl-CpG-binding domains to enrich for methylated DNA at sites of interest [10] and those based on sodium bisulfite treatment. However, the first three methods allow methylation detection over measured regions ranging in size from 100 to 1000 bp. Methods that use sodium bisulfite treatment, which converts unmethylated cytosines to thymine (via uracil) while methylated cytosines remain protected, measure DNA methylation at single nucleotide resolution [11]. For the remainder of this section, we will focus on bisulfite-conversion based sequencing techniques.

Whole genome bisulfite sequencing (WGBS) is considered the ‘gold standard’ for assaying DNA methylation because it provides global coverage at single-base resolution. Briefly, it combines bisulfite conversion of DNA molecules with high-throughput sequencing. To perform WGBS, the genomic DNA is first randomly fragmented to the desired size (200 bp). The fragmented DNA is converted into a sequencing library by ligation to adaptors that contain 5mCs. The sequence library is then treated with bisulfite. This treatment effectively converts unmethylated cytosines to uracil. After amplifying the library treated with bisulfite by PCR, it is sequenced using high-throughput sequencing. After the PCR, uracils will be represented as thymines. A precise recall of cytosine methylation requires not only sufficient sequencing depth, but also strongly depends on the quality of bisulfite conversion and library amplification. The benefit of this shotgun approach is that it typically reaches coverage of over 90% of the CpGs in the human genome in unbiased representation. It allows identification of non-CG methylation as well as identification of partially methylated domains (PMDs, [2]), and regions at distal regulatory elements with low methylation (LMRs, [12]) and DNA methylation valleys (DMVs) in embryonic stem cells [13]. Despite its advantages, WGBS remains the most expensive technique and standard library prep requires relatively large quantities of DNA (100ng–5 ug); as such, it is usually not applied to large numbers of samples [14]. To achieve high sensitivity in detecting methylation differences between samples, high sequencing depth is required which leads to significant increase in sequencing cost.

Reduced representation bisulfite sequencing (RRBS) is another technique that can also profile DNA methylation at single-base resolution. It combines digestion of genomic DNA with restriction enzymes and sequencing with bisulfite treatment in order to enrich for areas with high CpG content. Thus, it relies first on digestion of genomic DNA with restriction enzymes, such as MspI which recognises 5’-CCGG-3’ sequences and cleaves the phosphodiester bonds upstream of CpG dinucleotide. It can sequence only CpG dense regions and doesn’t interrogate CpG-deficient regions such as functional enhancers, intronic regions, intergenic regions or in general lowly methylated regions (LMRs) of the genome. It has limited coverage of the genome in CpG-poor regions and examines about 4% to 17% of the approximately 28 million CpG dinucleotides distributed throughout the human genome depending on the sequencing depth and which variant of RRBS is used [15,16].

Targeted Bisulfite sequencing also uses a combination of bisulfite sequencing with high-throughput sequencing, but it needs a prior selection of predefined genomic regions of interest. Frequently used protocols employ either PCR amplification of regions of interest [17,18], padlock probes [19], hybridization-based target enrichment [20], or convert-then-capture approaches [21].

One of the major assay specific issues is the fact that bisulfite sequencing cannot discriminate between hydroxymethylation (5hmC) and methylation (5mC) [22]. Hydroxymethylation converts to cyto-5-methanesulfonate upon bisulfite treatment, which then reads as a C when sequenced [22]. Furthermore, 5hmC mediated by TET proteins is a mechanism of non-passive DNA demethylation. Hence, methylation measurements for tissues having high 5-hydroxymethylation will be unreliable at least in certain genomic regions. The development of Tet-assisted bisulfite sequencing (TAB-seq) [23] and oxBS-Seq [24] has made it possible to distinguish between the two modifications with single-base resolution. In addition to 5hmC, single-base resolution mapping of 5caC using CAB-seq [25] and detection of 5fc (fCAB-seq [26,27] and redBS-Seq [26,27]) in mammalian genomes has recently been achieved.

## 3. Alignment and data processing for bisulfite sequencing

Since BS-seq changes unmethylated cytosines (C) to thymines (T), subsequent analysis steps focus on counting the number of C to T conversions and quantifying the methylation proportion per base. This is simply done by identifying C-to-T conversions in the aligned reads and dividing number of Cs by the sum of Ts and Cs for each cytosine in the genome. Being able to do the quantification reliably depends on quality control before alignment, the alignment methods and post-alignment quality control.

Since base-calling quality is not constant and could change between sequencing runs and within the same read, it is important to check the base quality (which represents the level of confidence in the base calls). Miscalled bases can be counted as C-T conversions erroneously, and such errors should be avoided if possible. This basic quality check can be done via fastQC software (http://www.bioinformatics.babraham.ac.uk/projects/fastqc/). Furthermore, sometimes adapters can be sequenced and if not properly removed, they will either lower the alignment rates or cause false C-T conversions. We recommend trimming low quality bases on sequence ends and removing adapters to minimize issues with false C-T conversions and to increase alignment rates. This can be achieved using trimming programs such as Trim Galore (http://www.bioinformatics.babraham.ac.uk/projects/trim_galore/).

Once pre-alignment quality control and processing is done, the next step is the alignment where potential C-T conversions should be handled. The BS-seq alignment methods mostly rely on modifications of known short-read alignment methods. For example, Bismark relies on Bowtie and in silico C-T conversion of reads and genomes [28]. Many other aligners use this *in silico* conversion strategy, such as: MethylCoder [29], BS-seeker2 [30], BRAT-BW [31] and Bison [32]. Other methods, such as Last [33], use a specific score matrix that can tolerate C-T mismatches or, such as BSMAP [34], masks Ts in the reads and matches them to genomic Cs. There are few comprehensive benchmarks of the aligners since new alternatives emerge frequently, but earlier attempts to compare the performance of the aligners did not find sufficient differences between aligners to exclude any from consideration[35,36]. Furthermore, recent tools are usually only better in some aspects of the benchmark; they may, for example, outperform competing tools in terms of computing time, but show a much higher memory footprint or have a worse mapping efficiency [35,36]. Some of these performance differences even disappear by varying parameters of the tools [37] and we see no compelling evidence that an established tool such as Bismark is significantly worse or better in accuracy than competing tools. For our own work, we frequently use Bismark since it provides BAM files, as well as additional methylation call related metrics and files.

After the alignment and methylation calling, there is still a need for further quality control. There are potential problems to be highlighted here. During the end repair step following the fragmentation unmethylated Cs are introduced at the ends of the DNA fragments [38]. This leads to a significant drop in the average methylation level that can be detected in a methylation bias (M-bias) plot [38,39] at those ends. A simple solution for this would be to disregard the affected positions in the sequenced reads [38,39]. Furthermore, incomplete conversion can occur during bisulfite treatment, where not all unmethylated Cs are converted to Ts [40]. Incomplete conversion causes false positive results due to interpretation of the unconverted unmethylated cytosines as methylated. For species without major non-CpG methylation, such as human, we can calibrate the conversion rate by using the percentage of non-CpG methylation. For a high quality experiment, we expect the conversion rate to be as close to 100% as possible, typical values for a good experiment will be higher than 99.5%. Another way to measure conversion rate is to add spike-in sequences with unmethylated Cs and counting the number of Ts for unmethylated Cs. Degradation of DNA during bisulfite treatment is another potential problem. Long incubation time and high bisulfite concentration, can lead to the degradation of about 90% of the incubated DNA [41]. Therefore, it is crucial to check unique alignment rates and read lengths after trimming. Moreover, it has been shown that the majority of CpGs with high inter-population differences contain common genomic SNPs (minor allele frequency > 0.01) [42]. To ensure more reliable interpretation of the data we advise removing known C/T SNPs which can interfere with methylation calls. The last post-alignment quality procedure addresses PCR bias. A simple way could be to remove reads that align to the exact same genomic position on the same strand. This de-duplication can be performed using the “samtools rmdup” command or Bismark tools. For RRBS, removing PCR duplicates by looking at overlapping coordinates of reads is not advised. Instead, one can try to remove PCR bias by removing regions with unusually high coverage; this method produces concurrent methylation measurements with orthogonal methods such as pyrosequencing [43].

## 4. Differential methylation methods

Once methylation proportions per base are obtained, generally, the dynamics of methylation profiles are considered next. When there are multiple sample groups, it is usually of interest to locate bases or regions with different methylation proportions across samples. The bases or regions with different methylation proportions across samples are called differentially methylated CpG sites (DMCs) and differentially methylated regions (DMRs). They have been shown to play a role in many different diseases due to their association with epigenetic control of gene regulation. In addition, DNA methylation profiles can be highly tissue-specific due to their role in gene regulation [44]. DNA methylation is highly informative when studying normal and diseased cells, because it can also act as a biomarker [44]. For example, the presence of large-scale abnormally methylated genomic regions is a hallmark feature of many types of cancers [45]. Because of aforementioned reasons, investigating differential methylation is usually one of the primary goals of doing bisulfite sequencing.

Differential DNA methylation is usually calculated by comparing the proportion of methylated Cs in a test sample relative to a control. In simple comparisons between such pairs of samples (i.e. test and control), methods such as Fisher’s Exact Test (implemented in e.g. methylkit [46] and RnBeads [47]) can be applied when there are no replicates for test and control cases. There are also methods based on hidden Markov models (HMMs) such as ComMet, included in the Bisulfighter methylation analysis suite [48,49] or the MethPipe software package [50]. These tools are sufficient to compare one test and one control sample at a time; if there are replicates, replicates can be pooled within groups to a single sample per group [46]. This strategy, however, does not take into account biological variability between replicates.

Regression-based methods are generally used to model methylation levels in relation to the sample groups and variation between replicates. Differences between currently available regression methods stem from the choice of distribution to model the data and the variation associated with it. In the simplest case, linear regression can be used to model methylation per given CpG or loci across sample groups. The model fits regression coefficients to model the expected methylation proportion values for each CpG site across sample groups. Hence, the null hypothesis of the model coefficients being zero could be tested using t-statistics. Such models are available in the limma package [51]. Limma was initially developed for the detection of differential gene expression in microarray data, but it is also used for methylation data. It is the default method applied in RnBeads. It uses moderated t-statistics in which standard errors have been moderated across loci, i.e. shrunk towards a common value using Empirical Bayes method. Another method that relies on linear regression and t-tests is the BSmooth [39] method. The main difference is that BSmooth applies a local-likelihood smoother to smooth DNA methylation across CpGs within genomic windows, assumes that data follow a binomial distribution and parameters are estimated by fitting linear model inside windows. It calculates signal-to-noise ratio statistic similar to t-test together with Empirical Bayes approach to test the difference for each CpG.

However, linear regression based methods might produce fitted methylation levels outside the range [0, 1] unless the values are transformed before regression. An alternative is logistic regression, which can deal with data strictly bounded between 0 and 1 and with non-constant variance, such as methylation proportion/fraction values. In the logistic regression, it is assumed that fitted values have variation *np(1-p)*, where *p* is the fitted methylation proportion for a given sample and *n* is the read coverage. If the observed variance is larger or smaller than assumed by the model, one speaks of under- or overdispersion. This over/under-dispersion can be corrected by calculating a scaling factor and using that factor to adjust the variance estimates as in *np(1-p)s,* where *s* is the scaling factor. MethylKit can apply logistic regression to test the methylation difference with or without the overdispersion correction. In this case, Chi-square or F-test can be used to compare the difference in the deviances of the null model and the alternative model. The null model assumes there is no relationship between sample groups and methylation, and the alternative model assumes that there is a relationship where sample groups are predictive of methylation values for a given CpG or region for which the model is constructed.

More complex regression models use beta binomial distribution and are particularly useful for better modeling the variance. Similar to logistic regression, their observation follows binomial distribution (number of reads), but methylation proportion itself can vary across samples, according to a beta distribution. It can deal with fitting values in [0,1] range and performs better when there is greater variance than expected by the simple logistic model. In essence, these models have a different way of calculating a scaling factor when there is overdispersion in the model. Further enhancements are made to these models by using the Empirical Bayes methods that can better estimate hyperparameters of beta distribution (variance-related parameters) by borrowing information between loci or regions within the genome to aid with inference about each individual loci or region. Some of the tools that rely on beta-binomial or beta model are as follows: MOABS [52] and DSS [53], RADMeth [54], BiSeq [52,55] and methylSig [56].

The choice of which method to apply also depends on the data at hand. If replicates are not available, possible tests include Fisher’s Exact test (implemented in methylKit, RnBeads and along with many other tool) or HMM-based methods such as ComMet. If replicates are available, tests based on regression are the natural choice rather than pooling the sample groups. Regression methods also have the advantage that one can add covariates into the tests such as technical/batch effects effects, age, sex, cell type heterogeneity, and genetic effects. For instance, it has been shown that age is a contributing factor for methylation values at some CpGs [57,58] and genetic heritability [59]. Covariates can be added to many methods such as methylKit, DSS, BSmooth and RnBeads.

Various differential methylation detection tools are based on similar methods and each method has its own advantages and disadvantages. To show this, we compared three classes of methods: 1) t-test/linear regression, 2) logistic regression and 3) beta binomial regression. For comparisons, we used both a simulated data set and a biologically relevant data set where we expect differentially methylated bases in certain regions. For the simulated data set, we used three different tools: DSS (beta binomial regression), limma (linear regression), and methylKit (logistic regression with/without overdispersion correction). We simulated a dataset consisting of 6 samples (3 controls and 3 samples with treatment). The read coverage modeled by a binomial distribution. The methylation background followed a beta distribution with parameters alpha=0.4, beta=0.5 and theta=10. We simulated 6 sets of 5000 CpG sites where methylation at 50% of the sites was affected by the treatment to varying degrees - specifically, methylation was elevated by 5%, 10%, 15%, 20% and 25% with respect to the test sample respectively. To adjust p-values for multiple testing, we used the q-value method [60] and we defined differentially methylated CpG sites with q-values below 0.01 for all examined methods. We calculated sensitivity, specificity and F-score for each of the three methods above. Sensitivity measured the proportion of true differentially methylated CpGs that were correctly identified as such, specificity was calculated as the proportion of detected CpGs that were truly not differentially methylated and correctly identified as such and F-score refers to a way to measure sensitivity and specificity by calculating their harmonic mean. Limma detected the fewest DMCs and consequently the fewest true positives (see Suppl. Figure 1) which lead to the lowest sensitivity (Figure 2a). DSS had similar results to limma where both also had high specificity (Figure 2b). MethylKit also performed well using either the Chi-squared or F-test. MethylKit without overdispersion showed the lowest specificity (the overdispersion correction usually improves specificity). F-test with overdispersion has similar results to DSS, whereas the Chi-squared test with overdispersion correction has similar specificity to stringent methods such as DSS and limma but achieves higher sensitivity (Figure 2c). Overall, DSS and limma are not very sensitive but very specific. We believe that a good compromise between the DSS/limma/F-test and default methylKit test is the overdispersion corrected methylKit Chi-square test. In addition, higher effect sizes results in higher number of detected true positives, higher sensitivity for all methods, and higher number of DMCs detected jointly by all methods (Suppl. Figure 2). Researchers should also consider a cutoff for the effect size or methylation difference in their analyses, as it is easier to detect changes with higher effect sizes and smaller effect sizes may not be biologically meaningful. A 5% change in methylation may not have an equivalent effect on gene expression and small changes may be within the range of the acceptable noise for biological systems.

**Figure 2.**
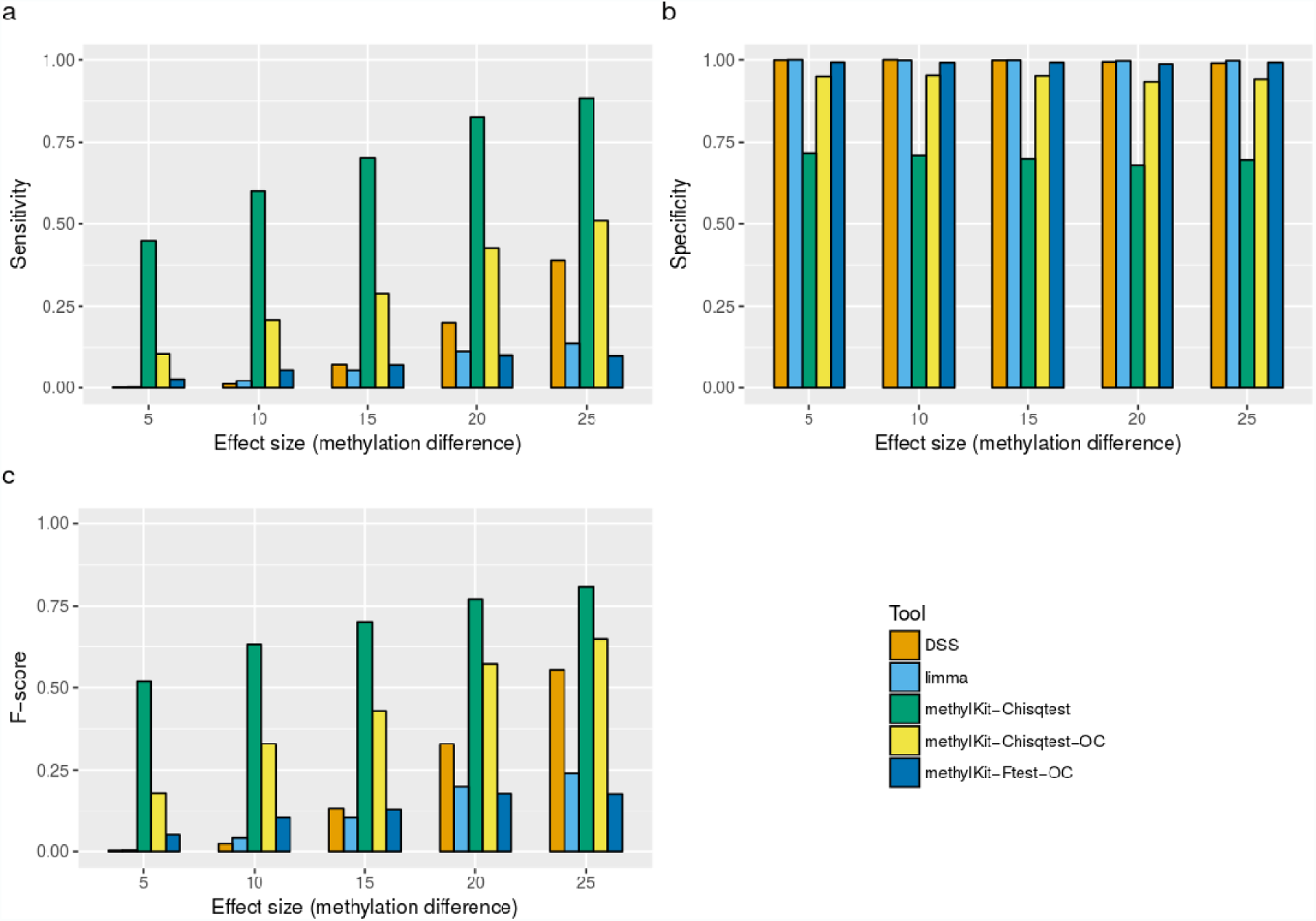
Comparison of DMC detection methods on simulated data. Barplots show sensitivity (a), specificity (b) and F-score (c) using DSS, limma, methylKit with Chi-squared or F-test. Overdispersion correction available only for methylKit has a suffix “-OC”. Effect size indicates methylation differences between two groups of samples (treatment and normal samples). Replicates in one group had elevated methylation in 50% of CpGs sites by accordingly 5%, 10%, 15%, 20% and 25%.

The performance of different methods using simulated datasets are always a subject of debate. There are many different ways to simulate datasets and how the data is simulated can bias the performance metrics towards certain methods. Therefore, we also compared the performance of different methods using real bisulfite sequencing experiments where we expect to see changes between samples in certain locations. Stadler and colleagues showed that DNA-binding factors can create low-methylated regions upon binding [12]. One of them, a CTCF protein, is a TF CCCTC-binding factor (zinc finger protein) that has a critical role in complex genome processes such as transcription, long range interactions, subnuclear organisation [61] and imprinting [62]. It had been shown that reduced methylation is a feature of CTCF-occupied sites supported by a high CpG content and specific CTCF recognition sequences, and if the site is unoccupied, the region on and around the site will have high methylation [63]. This means that if the CTCF occupancy changes between two cell types, we expect to see a change in the methylation levels as well. With this information, we looked for differentially methylated bases in regions that gained or lost CTCF binding between two cell types. We used the CTCF occupancy peaks supported by a presence of CTCF DNA motifs derived from the Factorbook database, binarized as ‘peak present’ or ‘peak lost’, and the ENCODE RRBS data (where each cell line has two replicates) for 19 human cell lines [64]. We performed pairwise comparisons for each pair in all possible combinations of these 19 cell lines. We defined true positives as the number of CTCF peaks gained/lost between two cell lines which overlap with at least one DMC. True negatives are defined as the number of CTCF peaks that do not change between cell lines and do not overlap any DMC even though they are covered by RRBS reads. Accordingly, false positives are defined as the number of CTCF peaks that are present in both cell lines but overlap with at least one DMC, while false negatives are defined as peaks that are gained or lost between cell lines but have no DMC. We also down-sampled the CTCF peaks that do not change to match the number of peaks that change, in order to have a balanced classification performance. Without this correction, true negatives overwhelm performance metrics since there are many CTCF peaks that do not change. Differentially methylated CpGs were identified for all combinations of two cell lines using DSS, limma, methylKit and BSmooth. In the simulation data set, we did not model changes in methylation of nearby CpGs and since BSmooth assumes that the true methylation profile is smooth and uses a local smoother, it was not adequate to apply this method on simulation data and did not perform well.

For the CTCF dataset, we observed consistent results with the simulated dataset results (see Figure 3). limma has the highest specificity (Figure 3b), however it detects extremely small number of true positives (Supp. Figure 3) and has the lowest sensitivity (Figure 3a). MethylKit without overdispersion had the highest F-score (Figure 3c), but also the lowest specificity. With overdispersion, methylKit showed higher specificity close to DSS and BSmooth and second highest F-score. methylKit and DSS show similar methylation level of true DMCs (Figure 3d). limma can only capture CpGs with higher methylation difference/effect size and BSmooth has the lowest methylation differences due to the smoothing step performed before computing the t-statistics. Taken together with the simulation results, methylKit without overdispersion can be used for more exploratory analysis as it achieves higher sensitivity but lower specificity, although it is still the best method when overall accuracy is considered. In contrast limma, DSS and methylKit F-test with overdispersion correction can be applied when there is a need to limit false positive rates, such as when picking regions or CpGs for validation. A good compromise between stringent and relaxed methods seems to be Chi-squared test with overdispersion correction.

**Figure 3.**
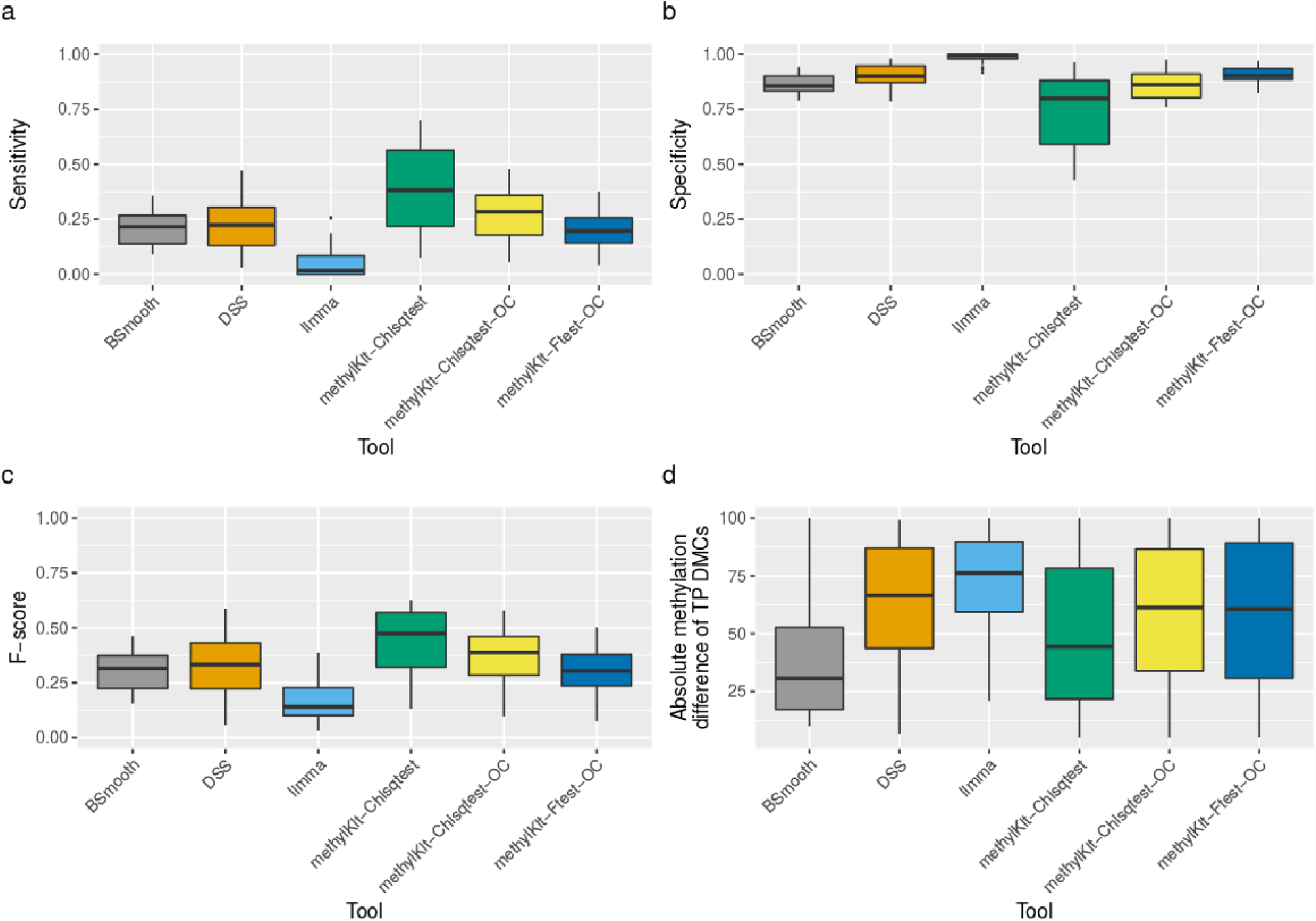
Performance measurements of tools for DMCs detection based on the association between CTCF occupancy with methylation status in cell-type specific manner using the Wang et al data and the RRBS ENCODE data. Barplots show sensitivity (a), specificity (b) and F-score (c) using BSmooth, DSS, limma, methylKit between pairs of multiple cell lines. MethylKit was performed using Chi-squared and F-test. MethylKit with overdispersion correction is depicted with “-OC” suffix. The absolute methylation percentage differences of DMCs found in CTCF peaks, that for given two cell lines, in one cell line has a gain and another lost of occupancy (true positives) are shown in subfigure d.

### 4.1 Defining differentially methylated regions

Most of the methods for differential methylation calling discussed earlier are designed to calculate both DMCs and DMRs. Some of them are designed to detect DMRs via aggregating DMCs together within a predefined regions, such as CpG islands or CpG shores. RADmeth [54] and eDMR [65] group P-values of adjacent CpGs and produce differentially methylated regions based on distance between differential CpGs and combination of their P-values using weighted Z-test. DSS set some thresholds on the P-values, number of CpG sites and length of regions before aggregation. Similarly, BSmooth defines DMRs by taking consecutive CpGs and cutoff based on the marginal empirical distribution of t and DMRs are ranked by sum of t-statistics in each CpG. BiSeq, on the other hand, first agglomerates CpG sites into clusters and smoothes methylation within clusters, uses beta regression and Wald test to test a group effect between control and test samples (with maximum likelihood with bias reduction). Apart from the various ways of clustering nearby CpGs or DMCs, many other methods rely on HMMs or other segmentation methods to segment the differential CpGs into hypo- and hyper-methylated regions and combine them to DMRs, such as MOABS, Methpipe, ComMet and methylKit.

Other methods define DMRs directly based on pre-defined windows. When input for functions for differential methylation calling are regions, so then data is summarized per region. The regions can be either predefined (such as regions with biological meaning like CpG islands) or user-defined with criteria like fixed region length for tiling windows that cover the whole genome, fixed numbers of significant adjacent CpG sites and smoothed estimated effect sizes.

## 5. Segmentation of the methylome

The analysis of methylation dynamics is not exclusively restricted to differentially methylated regions across samples, apart from this there is also an interest in examining the methylation profiles within the same sample. Usually, depressions in methylation profiles pinpoint regulatory regions like gene promoters that co-localize with CG-dense CpG islands. On the other hand, many gene-body regions are extensively methylated and CpG-poor [1]. These observations would describe a bimodal model of either hyper- or hypomethylated regions dependent on the local density of CpGs [66]. However, given the detection of CpG-poor regions with locally reduced levels of methylation (on average 30 %) in pluripotent embryonic stem cells and in neuronal progenitors in both mouse and human, a different model seems also reasonable [12]. These low-methylated regions (LMRs) are located distal to promoters, have little overlap with CpG islands and associated with enhancer marks such as p300 binding sites and H3K27ac enrichment.

The identification of these LMRs can be achieved by segmentation of the methylome using computational approaches. One of the well-known segmentation methods is based on a three-state Hidden Markov Model (HMM) taking only DNA methylation into account, without knowledge of any additional genomic information such as CpG density or functional annotations [12]. The three states that the authors aimed for were fully methylated regions (FMRs), unmethylated regions (UMRs) and low-methylated regions (LMRs). This segmentation represents a summary of methylome properties and features, in which unmethylated CpG islands correspond to UMRs [5], the majority is classified as FMR since most of the genome is methylated [67] and LMRs represent a new feature with intermediate levels of methylation, poor CpG content and shorter length compared to CpG islands [12]. Other segmentation methods such as MethPipe assume a two model state HMM and can not differentiate between LMRs and UMRs.

The authors of the R package “MethylSeekR” [68] adapt the idea of a three-state methylome and additionally identify partially methylated domains (PMDs), another methylome feature found, for instance, in human fibroblast but not in H1 embryonic stem cells [2,69]. These large regions, spanning hundreds of kilobases, are characterized by highly disordered methylation with average levels of methylation below 70% and covering almost 40% of the genome [2,69]. PMDs do not necessarily occur in every methylome, but their presence can be detected using a sliding window statistic [68]. In both MethylSeekR and MethPipe, the genome wide identification is done by training a two-state HMM, to separate PMDs from background regions. Then the PMDs are masked prior the characterization of UMRs/LMRs or hyper-/hypomethylated regions [50],[68].

There are also other segmentation strategies based on change-point analysis, where change-points of a genome-wide signal are recorded and the genome is partitioned into regions between consecutive change points. This approach is typically used in the context of copy number variation detection [70] but can be applied to methylome segmentation as well. A package implementing this method of segmentation based on change points is methylKit. It identifies segments that are further clustered using a mixture modeling approach. This clustering is based on only the average methylation level of the segments and allows the detection of distinct methylome features comparable to UMRs, LMRs and FMRs. This approach provides a more robust approach to segmentation where one can decide on the number of segmentation classes after segmentation. Whereas in HMM-based methods, one must know, *a priori,* the number of segmentation classes or run multiple rounds of HMMs with different numbers and identify which model fits best to the data.

### 5.1 Comparison of segmentation methods

We compared the change-point based segmentation to MethylSeekR, the latter of which is partially based on HMMs but mainly using cutoffs for methylation values. We identified high-concordance between these two methods by analysing chromosome 2 of the H1 embryonic stem cell methylome from the Roadmap Epigenomics Project [71]. They describe regions with similar segment lengths, number of CpGs per segment, methylation values and genome annotation (Figure 4a-d, respectively).

**Figure 4.**
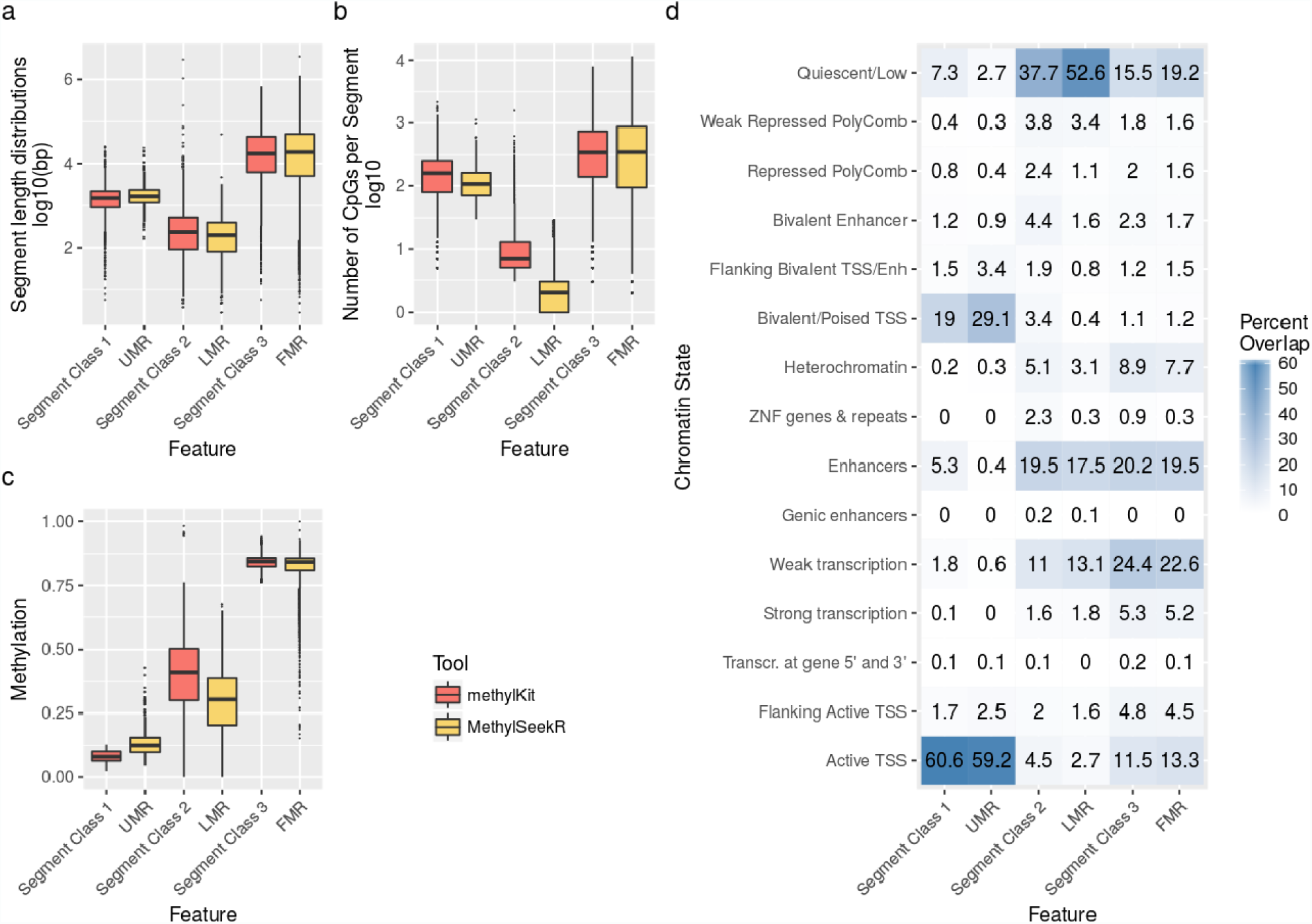
Comparison of features identified by segmentation tools analysing chromosome 2 of the H1 embryonic stem cells methylome. Boxplots show for each feature the distribution of (a) segment lengths in log10 transformed base pairs (bp) (b) CpG position covered by each segment in log10 transformed numbers (c) average methylation score per segment. (a) - (c) Boxplot colors indicate the tool generating the features either methylKit or MethylSeekR. (d) Heatmap showing the percentage of methylSeeker and methylKit segments that overlapped with chromatin state annotations for H1 embryonic stem cells.

We also applied change-point based segmentation to a genome with PMDs. We segmented the Human IMR90 methylome from the Roadmap Epigenomics Project [71] into four distinct features using methylKit. Then we compared feature-specific properties to published PMDs identified with MethPipe [50],[2,69] (Figure 5a-c) and found the feature with mean methylation level of segments closest to 50% to be the most proximate. We overlapped the published regions with all segments of this feature and found that 94% of the published regions of PMDs overlap with the generated segments of our feature (Figure 5d). In summary, change-point-based methods can be useful in the segmentation of the methylome, they provide classifications comparable to HMMs and also identify PMDs.

**Figure 5.**
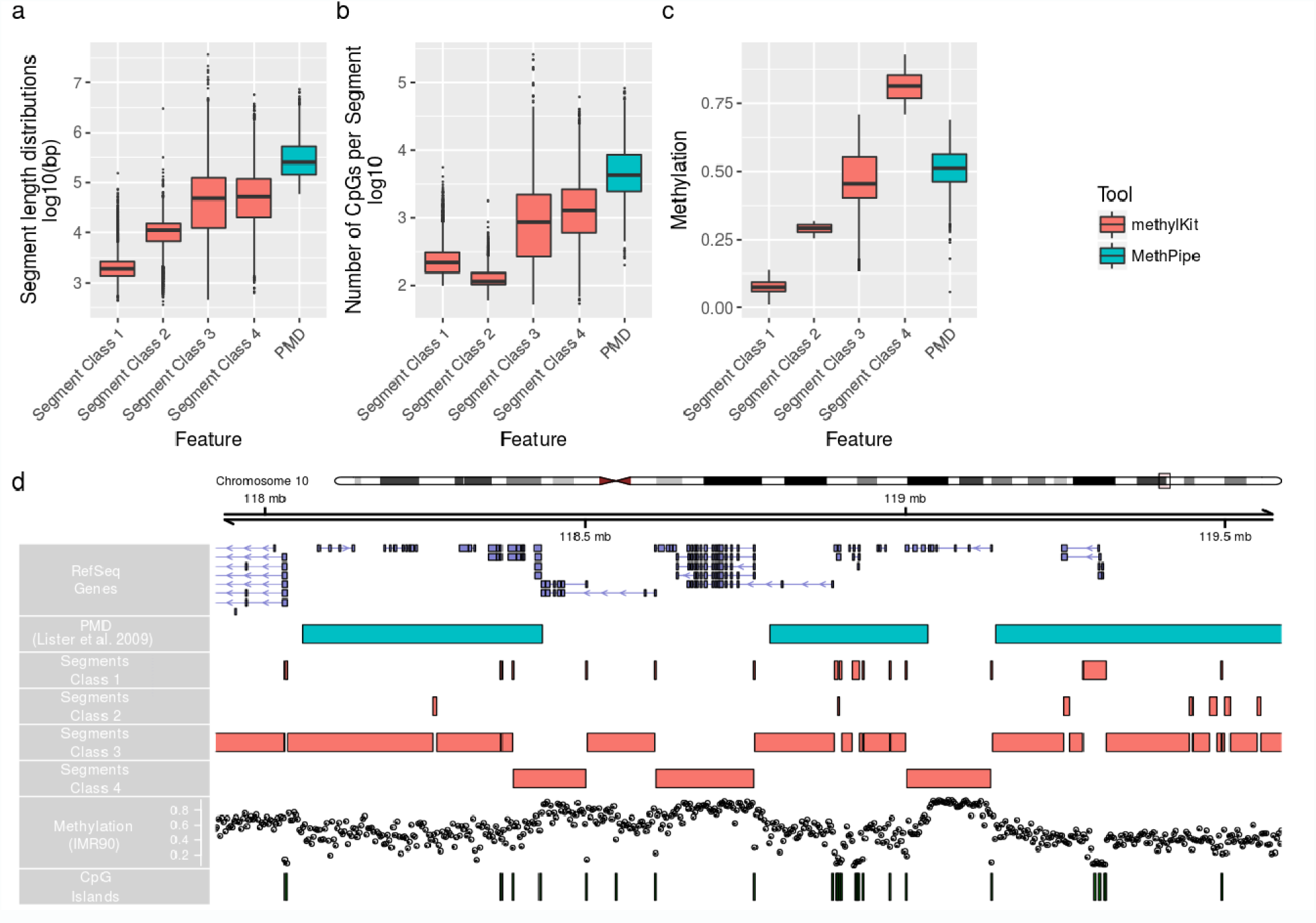
Comparison of features identified using methylKit change-point based segmentation on Human IMR90 methylome with published PMDs identified with MethPipe [50],[2,69]. Boxplots show for each feature the distribution of (a) segment lengths in log10 transformed base pairs (bp) (b) CpG position covered by each segment in log10 transformed numbers (c) average methylation score per segment. (a) - (c) Boxplot colors indicate the tool generating the features either methylKit or MethylPipe. (d) Genome Browser View showing region chr10: 117,786,314 - 121,271,788 of the hg19 assembly. The tracks are from top to bottom: “RefSeq Genes”, “Human_IMR90_PMD” from the Public “DNA Methylation” Track Hub [2],[50], Custom Segmentation Feature Track generated with methylKit, Human IMR90 methylome from Roadmap Epigenomics Project [71], CpG-Islands (track “cpgIslandExt”) from UCSC

## 6. Strategies for dealing with large datasets

With rising numbers of publicly available epigenetic data, it is tempting to reconstruct the results of published papers for many reasons, e.g. to better understand the reasoning behind steps the authors took or to develope a general intuition for the data. In the case of bisulfite sequencing data, we might want to perform differential methylation analysis in R using whole genome methylation data of multiple samples. The problem is that for genome-wide experiments, file sizes can easily range from hundreds of megabytes to gigabytes and processing multiple instances of those files in memory (RAM) might become infeasible unless we have access to a high performance cluster (HPC) with extensive RAM. If we want to use a desktop computer or laptop with limited RAM, we either need to restrict our analysis to a subset of the data or use packages that can handle this situation.

The authors of the RADmeth package for differential methylation analysis advise running the software on a “computing cluster with a few hundred available nodes” to allow the processing of multiple WGBS samples in a reasonable time. The same analysis can also be performed on a personal workstation with the disadvantage of increasing the computation time, which is in general dependent on three factors: the sample coverage, the number of sites analyzed and the number of samples. There exists one avenue to speed up the time-consuming step of regression if one’s workstation is a multicore system. The authors included a script to split the input data into smaller pieces which could than be processed separately and merged afterwards using UNIX commands.

A package for the comprehensive analysis of genome-wide DNA methylation data that can handle large data is RnBeads [47], which internally relies on the ‘ff’ package. The R package ‘ff’ [72] allows work with datasets larger than available RAM by storing them as temporary files and providing an interface to enable reading and writing from flat files and operate on the parts that have been loaded into R.

The methylKit package provides very similar capability by exploiting flat file databases to substitute in-memory objects if the objects grow too large. The internal data apart from meta information has a tabular structure storing chromosome, start/end position, strand information of the associated CpG base just like many other biological formats like BED, GFF or SAM. By exporting this tabular data into a TAB-delimited file and making sure it is accordingly position-sorted it can be indexed using the generic Tabix tool [66]. In general “Tabix indexing is a generalization of BAM indexing for generic TAB-delimited files. It inherits all the advantages of BAM indexing, including data compression and efficient random access in terms of few seek function calls per query.” [73]. MethylKit relies on Rsamtools (http://bioconductor.org/packages/release/bioc/html/Rsamtools.html) which implements tabix functionality for R and this way internal methylKit objects can be efficiently stored as compressed file on the disk and still be fast accessed. Another advantage is that existing compressed files can be loaded in interactive sessions, allowing the backup and transfer of intermediate analysis results.

## 7. Annotation of DMRs/DMCs and segments

The regions of interest obtained through differential methylation or segmentation analysis often need to be integrated with genome annotation datasets. Without this type of integration, differential methylation or segmentation results will be hard to interpret in biological terms. The most common annotation task is to see where regions of interest land in relation to genes and gene parts and regulatory regions: Do they mostly occupy promoter, intronic or exonic regions ? Do they overlap with repeats ? Do they overlap with other epigenomic markers or long-range regulatory regions ? These questions are not specific to methylation -nearly all regions of interest obtained via genome-wide studies have to deal with such questions. Thus, there are already multiple software tools that can produce such annotations. One is the Bioconductor package genomation [74]. It can be used to annotate DMRs/DMCs and it can also be used to integrate methylation proportions over the genome with other quantitative information and produce meta-gene plots or heatmaps. Another similar package is ChIPpeakAnno [75], which is designed for ChIP-seq peak annotation but could also be used for DMR/DMC annotation to a certain degree.

## 8. Workflows and tools that do not require programming experience

Software packages for the analysis of whole genome bisulfite sequencing data perform computationally intensive tasks and are therefore hosted on advanced hardware infrastructures. Moreover, the majority of the tools require programming knowledge (e.g. writing R commands). If the local execution of those tools is not feasible due to insufficient processing power or expertise, using an online service could be an alternative. For example, an analysis workflow on the RnBeads web service is started by simply uploading the data and setting a handful of options through a web form. The limitations it imposes on file size, however, make it infeasible for large datasets. Galaxy is an open source, web-based platform for data intensive biomedical research (see https://galaxyproject.org), providing access to publicly available servers and tools dedicated to data processing and analysis. A curated list of tools exists at https://toolshed.g2.bx.psu.edu hosting 4300 different programs for use within Galaxy at the time of writing, including methylKit https://toolshed.g2.bx.psu.edu/view/rnateam/methylkit/a8705df7c57f) and RnBeads (https://toolshed.g2.bx.psu.edu/view/pavlo-lutsik/rnbeads/6b0981ab063e). WBSA is another freely available^1^ web service for WGBS and RRBS (http://wbsa.big.ac.cn/) data. It is a modular collection of custom scripts combined with widely used tools, such as BWA for alignment and FastQC for quality control. The focus of WBSA is on ease of use. Uploading data and setting up analysis parameters is achieved using a small web form. The main advantages of this service are support for genome assemblies from 10 species, support for a range of sequencing protocols, as well as extraction and analysis of non-CpG methylation. More flexibility can be achieved by downloading and locally installing the modules, however, installing the WBSA back-end is a non-trivial task as its long list of dependencies includes tools and libraries from heterogeneous platforms: Java, MySQL, Perl, and R.

## 9. Conclusions

In this review, we have discussed the experimental and the computational methods for measuring and analysing DNA methylation in a genome-wide or targeted manner. We presented all the necessary steps of downstream analysis for bisulfite sequencing experiments starting from read alignment and quality check. We discussed and compared differential methylation and methylome segmentation methods. Our efforts for comparing differential methylation methods revealed that performances of different methods are comparable. One can choose methods based on the overall goal of their research. The methods that are stringent and limit the false positive rates are good for subsequent validation studies (DSS, limma, BSmooth, MethylKit with F-test and overdispersion correction), however these methods sacrifice sensitivity (true positive rate) for the sake of reducing false positives. A very relaxed method, such as the default methylKit method, has the best accuracy overall but highest false positive rate. A good alternative to stringent and relaxed methods is Chi-square test after overdispersion correction (implemented in methylKit). This method has high sensitivity without sacrificing too much for specificity. For segmentation methods, we observed high-concordance between cutoff-based methods and change-point analysis based methods. Change-point analysis methods are more flexible in the sense that they identify multiple biologically relevant segments within the same analysis. For example, HMM or cutoff based methods should first remove partially methylated domains (PMDs) from the analysis in order to define LMRs. Whereas methods based on change-point analysis can identify LMRs and PMDs in the same step.

We believe with this guideline of methods for BS-seq analysis both bioinformaticians and experimental biologists will gain insight into experimental design as well as best practises for computational analysis. The code that we used to generate the results is available online on the website: https://github.com/BIMSBbioinfo/Strategies_for_analyzing_BS-seq

## Acknowledgements and financial support

The authors acknowledge support from the German Federal Ministry of Education and Research (BMBF) as part of the RBC, de.NBI-epi and HD-HuB services centers of the German Network for Bioinformatics Infrastructure (de.NBI). We also acknowledge support for KW from Berlin Institute of Health (BIH). Authors would like to thank Brendan Osberg for valuable comments on the manuscript.

## Supplementary figures

**Suppl. Figure 1.**
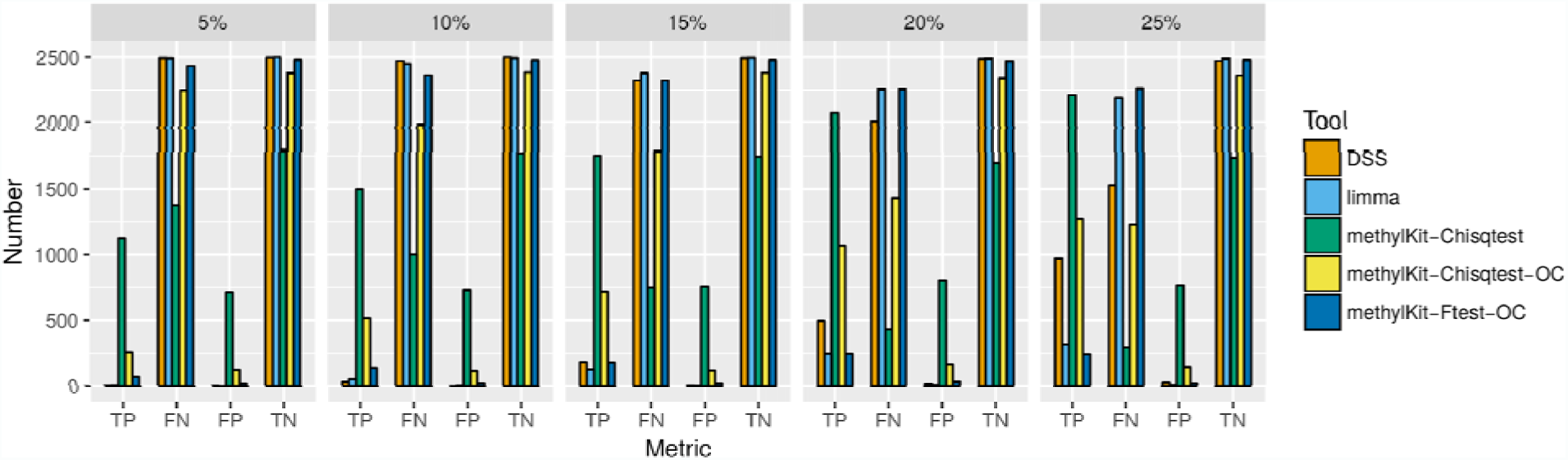
Barplots show number of true positives (TP), false negatives (FN), false positives (FP) and true negatives (TN) for each effect size (5%, 10%, 15%, 20% and 25%) on the simulated dataset using DSS, limma and methylKit tools.

**Suppl. Figure 2.**
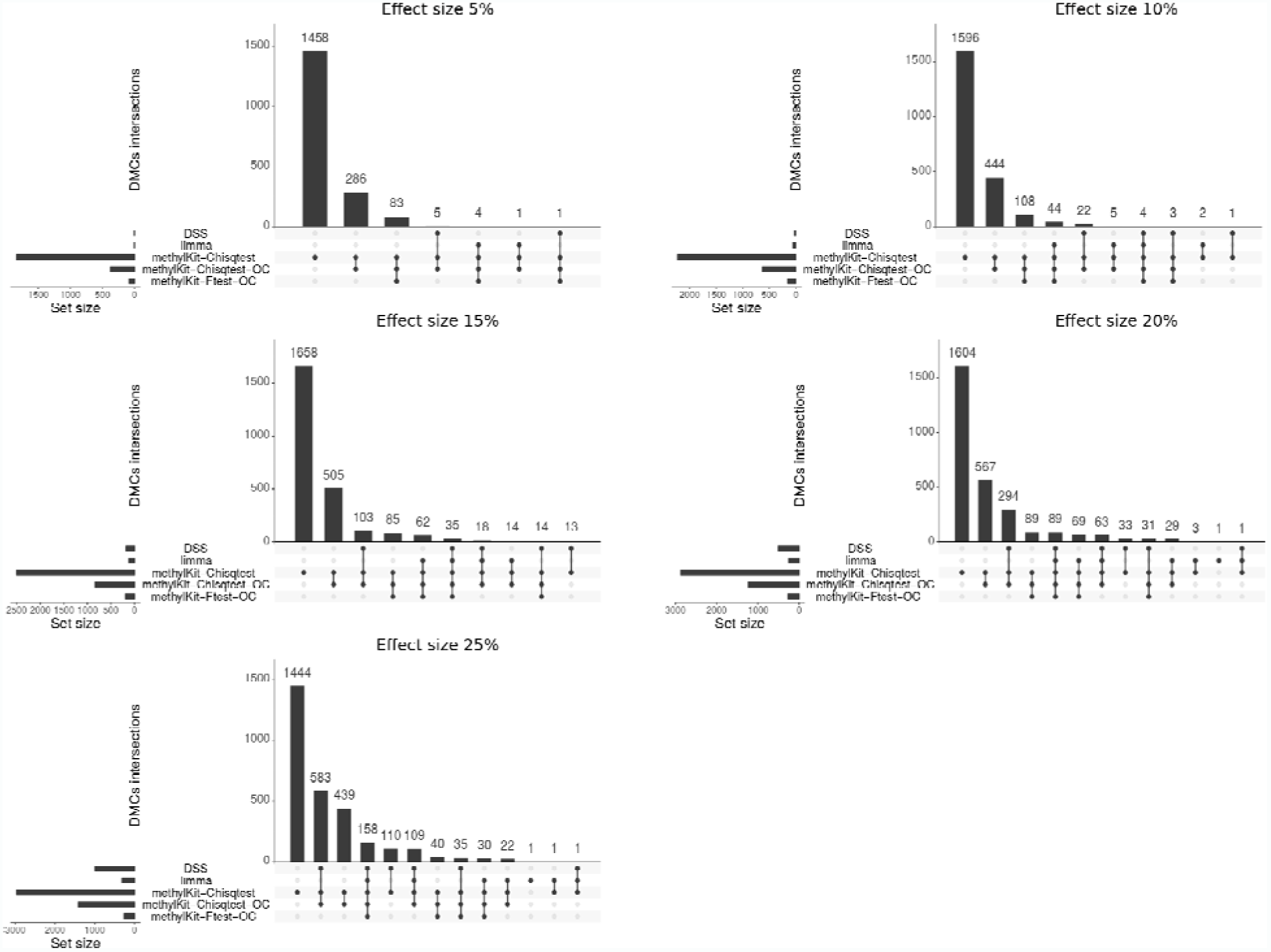
Visualisation of common and unique DMCs generated by tools: DSS, limma, methylKit using F-test with an overdispersion and Chisq test with and without the overdispersion correction using the simulated datasets. Empty intersections are now shown.

**Suppl. Figure 3.**
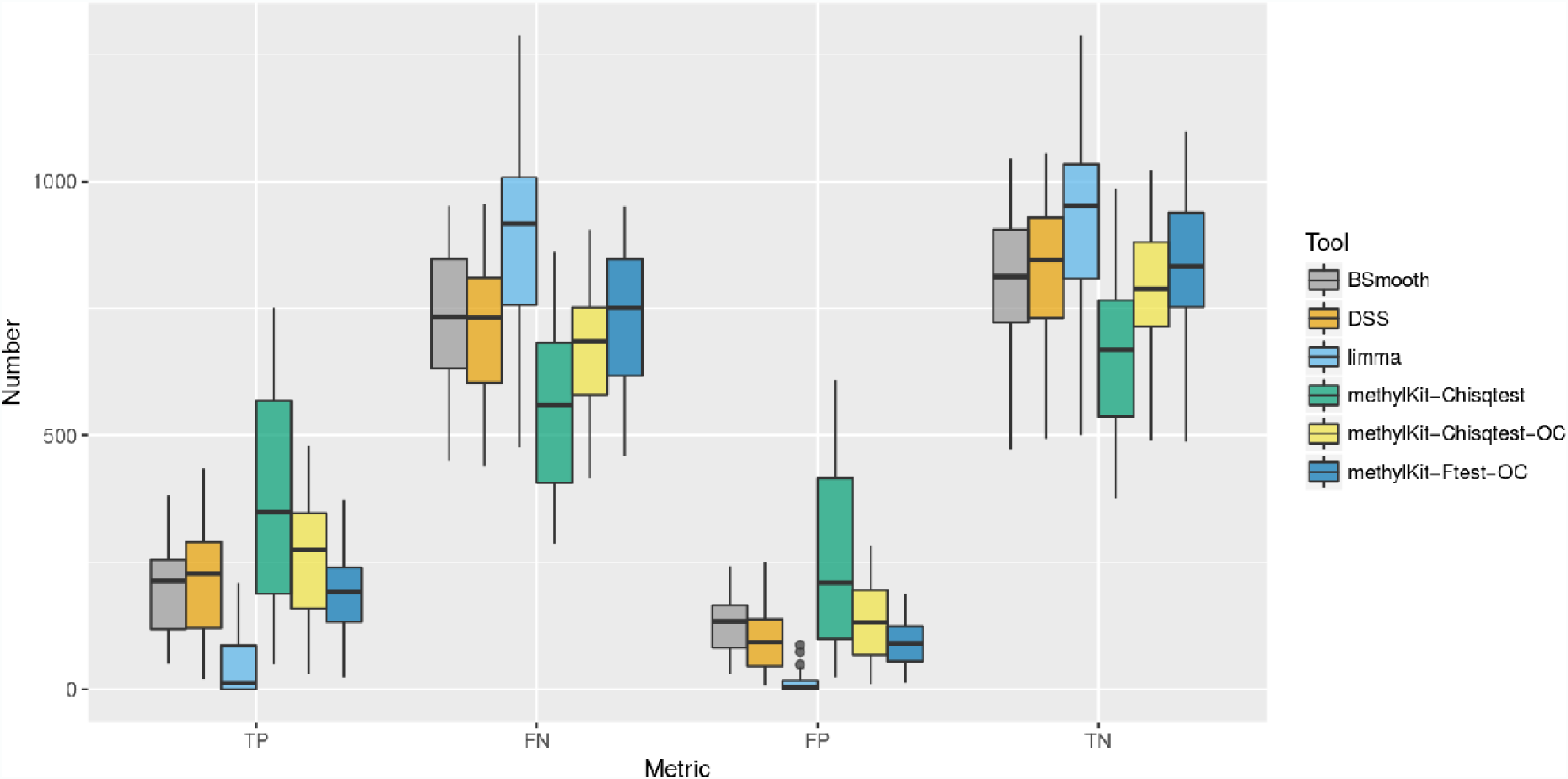
Number of true positives (TP), false negatives (FN), false positives (FP) and true negatives (TN) for each effect size (5%, 10%, 15%, 20% and 25%) on the CTCF occupancy with methylation status in cell-type specific manner using Wang et al data and the RRBs ENCODE data using DSS, limma and methylKit tools.

## Footnotes

1 for academic use only

## References

1. Bock C, Beerman I, Lien W-H, Smith ZD, Gu H, Boyle P, et al. DNA methylation dynamics during in vivo differentiation of blood and skin stem cells. Mol Cell. 2012;47: 633–647. doi:10.1016/j.molcel.2012.06.019

2. Lister R, Pelizzola M, Dowen RH, Hawkins RD, Hon G, Tonti-Filippini J, et al. Human DNA methylomes at base resolution show widespread epigenomic differences. Nature. 2009;462: 315–322. doi:10.1038/nature08514

3. Lister R, Mukamel EA, Nery JR, Urich M, Puddifoot CA, Johnson ND, et al. Global Epigenomic Reconfiguration During Mammalian Brain Development. Science. 2013;341: 1237905–1237905. doi:10.1126/science.1237905

4. Smith ZD, Meissner A. DNA methylation: roles in mammalian development. Nat Rev Genet. 2013;14: 204–220. doi:10.1038/nrg3354

5. Deaton AM, Bird A. CpG islands and the regulation of transcription. Genes Dev. 2011;25: 1010–1022. doi:10.1101/gad.2037511

6. Tahiliani M, Koh KP, Shen Y, Pastor WA, Bandukwala H, Brudno Y, et al. Conversion of 5-methylcytosine to 5-hydroxymethylcytosine in mammalian DNA by MLL partner TET1. Science. 2009;324: 930–935. doi:10.1126/science.1170116

7. He Y-F, Li B-Z, Li Z, Liu P, Wang Y, Tang Q, et al. Tet-mediated formation of 5-carboxylcytosine and its excision by TDG in mammalian DNA. Science. 2011;333: 1303–1307. doi:10.1126/science.1210944

8. Clark TA, Spittle KE, Turner SW, Korlach J. Direct detection and sequencing of damaged DNA bases. Genome Integr. 2011;2: 10. doi:10.1186/2041-9414-2-10

9. Weber M, Davies JJ, Wittig D, Oakeley EJ, Haase M, Lam WL, et al. Chromosome-wide and promoter-specific analyses identify sites of differential DNA methylation in normal and transformed human cells. Nat Genet. 2005;37: 853–862. doi:10.1038/ng1598

10. Brinkman AB, Simmer F, Ma K, Kaan A, Zhu J, Stunnenberg HG. Whole-genome DNA methylation profiling using MethylCap-seq. Methods. 2010;52: 232–236. doi:10.1016/j.ymeth.2010.06.012

11. Baubec T, Akalin A. Genome-Wide Analysis of DNA Methylation Patterns by High-Throughput Sequencing. Field Guidelines for Genetic Experimental Designs in High-Throughput Sequencing. 2016. pp. 197–221. doi:10.1007/978-3-319-31350-4_9

12. Stadler MB, Murr R, Burger L, Ivanek R, Lienert F, Schöler A, et al. DNA-binding factors shape the mouse methylome at distal regulatory regions. Nature. 2011;480: 490–495. doi:10.1038/nature10716

13. Xie W, Schultz MD, Lister R, Hou Z, Rajagopal N, Ray P, et al. Epigenomic analysis of multilineage differentiation of human embryonic stem cells. Cell. 2013;153: 1134–1148. doi:10.1016/j.cell.2013.04.022

14. Stirzaker C, Taberlay PC, Statham AL, Clark SJ. Mining cancer methylomes: prospects and challenges. Trends Genet. 2014;30: 75–84. doi:10.1016/j.tig.2013.11.004

15. Meissner A, Gnirke A, Bell GW, Ramsahoye B, Lander ES, Jaenisch R. Reduced representation bisulfite sequencing for comparative high-resolution DNA methylation analysis. Nucleic Acids Res. 2005;33: 5868–5877. doi:10.1093/nar/gki901

16. Rampal R, Alkalin A, Madzo J, Vasanthakumar A, Pronier E, Patel J, et al. DNA hydroxymethylation profiling reveals that WT1 mutations result in loss of TET2 function in acute myeloid leukemia. Cell Rep. 2014;9: 1841–1855. doi:10.1016/j.celrep.2014.11.004

17. Landan G, Cohen NM, Mukamel Z, Bar A, Molchadsky A, Brosh R, et al. Epigenetic polymorphism and the stochastic formation of differentially methylated regions in normal and cancerous tissues. Nat Genet. 2012;44: 1207–1214. doi:10.1038/ng.2442

18. Taylor KH, Kramer RS, Davis JW, Guo J, Duff DJ, Xu D, et al. Ultradeep bisulfite sequencing analysis of DNA methylation patterns in multiple gene promoters by 454 sequencing. Cancer Res. 2007;67: 8511–8518. doi:10.1158/0008-5472.CAN-07-1016

19. Ball MP, Li JB, Gao Y, Lee J-H, LeProust EM, Park I-H, et al. Targeted and genome-scale strategies reveal gene-body methylation signatures in human cells. Nat Biotechnol. 2009;27: 361–368. doi:10.1038/nbt.1533

20. Ivanov M, Kals M, Kacevska M, Metspalu A, Ingelman-Sundberg M, Milani L. In-solution hybrid capture of bisulfite-converted DNA for targeted bisulfite sequencing of 174 ADME genes. Nucleic Acids Res. 2013;41: e72. doi:10.1093/nar/gks1467

21. Li Q, Suzuki M, Wendt J, Patterson N, Eichten SR, Hermanson PJ, et al. Post-conversion targeted capture of modified cytosines in mammalian and plant genomes. Nucleic Acids Res. 2015;43: e81. doi:10.1093/nar/gkv244

22. Huang Y, Pastor WA, Shen Y, Tahiliani M, Liu DR, Rao A. The behaviour of 5-hydroxymethylcytosine in bisulfite sequencing. PLoS One. 2010;5: e8888. doi:10.1371/journal.pone.0008888

23. Yu M, Hon GC, Szulwach KE, Song C-X, Jin P, Ren B, et al. Tet-assisted bisulfite sequencing of 5-hydroxymethylcytosine. Nat Protoc. 2012;7: 2159–2170. doi:10.1038/nprot.2012.137

24. Booth MJ, Branco MR, Ficz G, Oxley D, Krueger F, Reik W, et al. Quantitative sequencing of 5-methylcytosine and 5-hydroxymethylcytosine at single-base resolution. Science. 2012;336: 934–937. doi:10.1126/science.1220671

25. Lu X, Song C-X, Szulwach K, Wang Z, Weidenbacher P, Jin P, et al. Chemical modification-assisted bisulfite sequencing (CAB-Seq) for 5-carboxylcytosine detection in DNA. J Am Chem Soc. 2013;135: 9315–9317. doi:10.1021/ja4044856

26. Song C-X, Szulwach KE, Dai Q, Fu Y, Mao S-Q, Lin L, et al. Genome-wide profiling of 5-formylcytosine reveals its roles in epigenetic priming. Cell. 2013;153: 678–691. doi:10.1016/j.cell.2013.04.001

27. Booth MJ, Marsico G, Bachman M, Beraldi D, Balasubramanian S. Quantitative sequencing of 5-formylcytosine in DNA at single-base resolution. Nat Chem. 2014;6: 435–440. doi:10.1038/nchem.1893

28. Krueger F, Andrews SR. Bismark: a flexible aligner and methylation caller for Bisulfite-Seq applications. Bioinformatics. 2011;27: 1571–1572. doi:10.1093/bioinformatics/btr167

29. Pedersen B, Hsieh T-F, Ibarra C, Fischer RL. MethylCoder: software pipeline for bisulfite-treated sequences. Bioinformatics. 2011;27: 2435–2436. doi:10.1093/bioinformatics/btr394

30. Guo W, Fiziev P, Yan W, Cokus S, Sun X, Zhang MQ, et al. BS-Seeker2: a versatile aligning pipeline for bisulfite sequencing data. BMC Genomics. 2013;14: 774. doi:10.1186/1471-2164-14-774

31. Harris EY, Ponts N, Le Roch KG, Lonardi S. BRAT-BW: efficient and accurate mapping of bisulfite-treated reads. Bioinformatics. 2012;28: 1795–1796. doi:10.1093/bioinformatics/bts264

32. Ryan DP, Ehninger D. Bison: bisulfite alignment on nodes of a cluster. BMC Bioinformatics. 2014;15: 337. doi:10.1186/1471-2105-15-337

33. Frith MC, Mori R, Asai K. A mostly traditional approach improves alignment of bisulfite-converted DNA. Nucleic Acids Res. 2012;40: e100. doi:10.1093/nar/gks275

34. Xi Y, Li W. BSMAP: whole genome bisulfite sequence MAPping program. BMC Bioinformatics. 2009;10: 232. doi:10.1186/1471-2105-10-232

35. Tsuji J, Weng Z. Evaluation of preprocessing, mapping and postprocessing algorithms for analyzing whole genome bisulfite sequencing data. Brief Bioinform. 2016;17: 938–952. doi:10.1093/bib/bbv103

36. Kunde-Ramamoorthy G, Coarfa C, Laritsky E, Kessler NJ, Harris RA, Xu M, et al. Comparison and quantitative verification of mapping algorithms for whole-genome bisulfite sequencing. Nucleic Acids Res. 2014;42: e43. doi:10.1093/nar/gkt1325

37. Tran H, Porter J, Sun M-A, Xie H, Zhang L. Objective and comprehensive evaluation of bisulfite short read mapping tools. Adv Bioinformatics. 2014;2014: 472045. doi:10.1155/2014/472045

38. Bock C. Analysing and interpreting DNA methylation data. Nat Rev Genet. 2012;13: 705–719. doi:10.1038/nrg3273

39. Hansen KD, Langmead B, Irizarry RA. BSmooth: from whole genome bisulfite sequencing reads to differentially methylated regions. Genome Biol. 2012;13: R83. doi:10.1186/gb-2012-13-10-r83

40. Genereux DP, Johnson WC, Burden AF, Stöger R, Laird CD. Errors in the bisulfite conversion of DNA: modulating inappropriate- and failed-conversion frequencies. Nucleic Acids Res. 2008;36: e150. doi:10.1093/nar/gkn691

41. Grunau C, Clark SJ, Rosenthal A. Bisulfite genomic sequencing: systematic investigation of critical experimental parameters. Nucleic Acids Res. 2001;29: E65–5. Available: https://www.ncbi.nlm.nih.gov/pubmed/11433041

42. Daca-Roszak P, Pfeifer A, Żebracka-Gala J, Rusinek D, Szybinska A, Jarzab B, et al. Impact of SNPs on methylation readouts by Illumina Infinium HumanMethylation450 BeadChip Array: implications for comparative population studies. BMC Genomics. 2015;16: 1003. doi:10.1186/s12864-015-2202-0

43. Akalin A, Garrett-Bakelman FE, Kormaksson M, Busuttil J, Zhang L, Khrebtukova I, et al. Base-pair resolution DNA methylation sequencing reveals profoundly divergent epigenetic landscapes in acute myeloid leukemia. PLoS Genet. 2012;8: e1002781. doi:10.1371/journal.pgen.1002781

44. Schübeler D. Function and information content of DNA methylation. Nature. 2015;517: 321–326. doi:10.1038/nature14192

45. Ehrlich M. DNA methylation in cancer: too much, but also too little. Oncogene. 2002;21: 5400–5413. doi:10.1038/sj.onc.1205651

46. Akalin A, Kormaksson M, Li S, Garrett-Bakelman FE, Figueroa ME, Melnick A, et al. methylKit: a comprehensive R package for the analysis of genome-wide DNA methylation profiles. Genome Biol. 2012;13: R87. doi:10.1186/gb-2012-13-10-r87

47. Assenov Y, Müller F, Lutsik P, Walter J, Lengauer T, Bock C. Comprehensive analysis of DNA methylation data with RnBeads. Nat Methods. 2014;11: 1138–1140. doi:10.1038/nmeth.3115

48. Saito Y, Mituyama T. Detection of differentially methylated regions from bisulfite-seq data by hidden Markov models incorporating genome-wide methylation level distributions. BMC Genomics. 2015;16 Suppl 12: S3. doi:10.1186/1471-2164-16-S12-S3

49. Saito Y, Tsuji J, Mituyama T. Bisulfighter: accurate detection of methylated cytosines and differentially methylated regions. Nucleic Acids Res. 2014;42: e45. doi:10.1093/nar/gkt1373

50. Song Q, Decato B, Hong EE, Zhou M, Fang F, Qu J, et al. A reference methylome database and analysis pipeline to facilitate integrative and comparative epigenomics. PLoS One. 2013;8: e81148. doi:10.1371/journal.pone.0081148

51. Ritchie ME, Phipson B, Wu D, Hu Y, Law CW, Shi W, et al. limma powers differential expression analyses for RNA-sequencing and microarray studies. Nucleic Acids Res. 2015;43: e47. doi:10.1093/nar/gkv007

52. Sun D, Xi Y, Rodriguez B, Park HJ, Tong P, Meong M, et al. MOABS: model based analysis of bisulfite sequencing data. Genome Biol. 2014;15: R38. doi:10.1186/gb-2014-15-2-r38

53. Feng H, Conneely KN, Wu H. A Bayesian hierarchical model to detect differentially methylated loci from single nucleotide resolution sequencing data. Nucleic Acids Res. 2014;42: e69. doi:10.1093/nar/gku154

54. Dolzhenko E, Smith AD. Using beta-binomial regression for high-precision differential methylation analysis in multifactor whole-genome bisulfite sequencing experiments. BMC Bioinformatics. 2014;15: 215. doi:10.1186/1471-2105-15-215

55. Hebestreit K, Dugas M, Klein H-U. Detection of significantly differentially methylated regions in targeted bisulfite sequencing data. Bioinformatics. 2013;29: 1647–1653. doi:10.1093/bioinformatics/btt263

56. Park Y, Figueroa ME, Rozek LS, Sartor MA. MethylSig: a whole genome DNA methylation analysis pipeline. Bioinformatics. 2014;30: 2414–2422. doi:10.1093/bioinformatics/btu339

57. Xu C, Qu H, Wang G, Xie B, Shi Y, Yang Y, et al. A novel strategy for forensic age prediction by DNA methylation and support vector regression model. Sci Rep. 2015;5: 17788. doi:10.1038/srep17788

58. Heyn H, Li N, Ferreira HJ, Moran S, Pisano DG, Gomez A, et al. Distinct DNA methylomes of newborns and centenarians. Proc Natl Acad Sci U S A. 2012;109: 10522–10527. doi:10.1073/pnas.1120658109

59. McRae AF, Powell JE, Henders AK, Bowdler L, Hemani G, Shah S, et al. Contribution of genetic variation to transgenerational inheritance of DNA methylation. Genome Biol. 2014;15: R73. doi:10.1186/gb-2014-15-5-r73

60. Storey JD, Tibshirani R. Statistical significance for genomewide studies. Proc Natl Acad Sci U S A. 2003;100: 9440–9445. doi:10.1073/pnas.1530509100

61. Bonev B, Cavalli G. Organization and function of the 3D genome. Nat Rev Genet. 2016;17: 772. doi:10.1038/nrg.2016.147

62. Kurukuti S, Tiwari VK, Tavoosidana G, Pugacheva E, Murrell A, Zhao Z, et al. CTCF binding at the H19 imprinting control region mediates maternally inherited higher-order chromatin conformation to restrict enhancer access to Igf2. Proc Natl Acad Sci U S A. 2006;103: 10684–10689. doi:10.1073/pnas.0600326103

63. Maurano MT, Wang H, John S, Shafer A, Canfield T, Lee K, et al. Role of DNA Methylation in Modulating Transcription Factor Occupancy. Cell Rep. 2015;12: 1184–1195. doi:10.1016/j.celrep.2015.07.024

64. Wang H, Maurano MT, Qu H, Varley KE, Gertz J, Pauli F, et al. Widespread plasticity in CTCF occupancy linked to DNA methylation. Genome Res. 2012;22: 1680–1688. doi:10.1101/gr.136101.111

65. Li S, Garrett-Bakelman FE, Akalin A, Zumbo P, Levine R, To BL, et al. An optimized algorithm for detecting and annotating regional differential methylation. BMC Bioinformatics. 2013;14 Suppl 5: S10. doi:10.1186/1471-2105-14-S5-S10

66. Lövkvist C, Dodd IB, Sneppen K, Haerter JO. DNA methylation in human epigenomes depends on local topology of CpG sites. Nucleic Acids Res. 2016;44: 5123–5132. doi:10.1093/nar/gkw124

67. Bird A. DNA methylation patterns and epigenetic memory. Genes Dev. 2002;16: 6–21. doi:10.1101/gad.947102

68. Burger L, Gaidatzis D, Schübeler D, Stadler MB. Identification of active regulatory regions from DNA methylation data. Nucleic Acids Res. 2013;41: e155. doi:10.1093/nar/gkt599

69. Gaidatzis D, Burger L, Murr R, Lerch A, Dessus-Babus S, Schübeler D, et al. DNA sequence explains seemingly disordered methylation levels in partially methylated domains of Mammalian genomes. PLoS Genet. 2014;10: e1004143. doi:10.1371/journal.pgen.1004143

70. Klambauer G, Schwarzbauer K, Mayr A, Clevert D-A, Mitterecker A, Bodenhofer U, et al. cn.MOPS: mixture of Poissons for discovering copy number variations in next-generation sequencing data with a low false discovery rate. Nucleic Acids Res. 2012;40: e69. doi:10.1093/nar/gks003

71. Roadmap Epigenomics Consortium, Kundaje A, Meuleman W, Ernst J, Bilenky M, Yen A, et al. Integrative analysis of 111 reference human epigenomes. Nature. 2015;518: 317–330. doi:10.1038/nature14248

72. Adler D, Gläser C, Nenadic O, Oehlschlägel J, Zucchini W. ff: Memory-efficient Storage of Large Data on Disk and Fast Access Functions. R package version. 2014; 2–2.

73. Li H. Tabix: fast retrieval of sequence features from generic TAB-delimited files. Bioinformatics. 2011;27: 718–719. doi:10.1093/bioinformatics/btq671

74. Akalin A, Franke V, Vlahoviček K, Mason CE, Schübeler D. genomation: a toolkit to summarize, annotate and visualize genomic intervals. Bioinformatics. 2015;31: 1127–1129. doi:10.1093/bioinformatics/btu775

75. Zhu LJ, Gazin C, Lawson ND, Pagès H, Lin SM, Lapointe DS, et al. ChIPpeakAnno: a Bioconductor package to annotate ChIP-seq and ChIP-chip data. BMC Bioinformatics. 2010;11: 237. doi:10.1186/1471-2105-11-237

